# A Computational and Biochemical Study of -1 Ribosomal Frameshifting in Human mRNAs

**DOI:** 10.1101/2021.04.23.441185

**Authors:** Xia Zhou, Xiaolan Huang, Zhihua Du

## Abstract

−1 programmed ribosomal frameshifting (−1 PRF) is a translational recoding mechanism used by many viral and cellular mRNAs. −1 PRF occurs at a heptanucleotide slippery sequence and is stimulated by a downstream RNA structure, most often in the form of a pseudoknot. The utilization of −1 PRF to produce proteins encoded by the −1 reading frame is wide-spread in RNA viruses, but relatively rare in cellular mRNAs. In human, only three such cases of −1 PRF events have been reported, all involving retroviral-like genes and protein products. To evaluate the extent of −1 PRF utilization in the human transcriptome, we have developed a computational scheme for identifying putative pseudoknot-dependent −1 PRF events and applied the method to a collection of 43,191 human mRNAs in the NCBI RefSeq database. In addition to the three reported cases, our study identified more than two dozen putative −1 PRF cases. The genes involved in these cases are genuine cellular genes without a viral origin. Moreover, in more than half of these cases, the frameshift site locates far upstream (>250 nt) from the stop codon of the 0 reading frame, which is nonviral-like. Using dual luciferase assays in HEK293T cells, we confirmed that the −1 PRF signals in the mRNAs of CDK5R2 and SEMA6C are functional in inducing efficient frameshifting. Our findings have significant implications in expanding the repertoire of the −1 PRF phenomenon and the protein*-*coding capacity of the human transcriptome.

## Background

Programmed −1 ribosomal frameshifting (−1 PRF) is a translational recoding mechanism in which a certain percentage of the elongating ribosomes slip back one nucleotide along the mRNA and continue translation in the −1 reading frame (1,2). Often, two cis-acting signals in the mRNA are required to induce efficient frameshifting: a heptanucleotide “slippery sequence” and a shortly downstream RNA structure. Based on known −1 PRF events, the slippery sequences seem to have a consensus of X XXY YYZ (the triplets represent the P site and A site codons in the zero reading frame, X can be any nucleotide, Y can be A or U, Z can be A, C or U). During −1 PRF, the P site and A site tRNAs on the XXY and YYZ codons shift back one nucleotide to the XXX and YYY codons in the −1 reading frame. Efficient frameshifting is often stimulated by an RNA structure located several nucleotides downstream of the slippery sequence. In most known cases of −1 PRF, the frameshift stimulator RNA structure is an H (hairpin) type pseudoknot (3-7). An H-type pseudoknot is formed when a stretch of nucleotides within the loop region of a hairpin basepairs with a complementary sequence outside that loop (8-10).

It is well known that many RNA viruses, including the etiological agents for AIDS (HIV-1: human immunodeficiency virus type-1) and COVID-19 (SARS-CoV2: severe acute respiratory syndrome coronavirus 2), utilize −1 PRF to express certain key viral proteins at a defined ratio (1,6,11,12). It was found that the levels of −1 PRF are maintained within a relatively narrow range, increasing or decreasing in the −1 PRF efficiency outside of that range significantly attenuates the production of infectious virions (13-16). Therefore, perturbation of −1 PRF efficiency may represent a viable strategy for the development of antiviral therapeutics against viral pathogens. Efforts have been made to exploit the viral −1 PRF signals, especially the frameshift-stimulatory structures, as putative drug targets (17-25). Small organic compounds, peptides, antisense oligonucleotides and peptide nucleic acids (PNAs) have been identified that can modulate −1 PRF induced by the frameshifting signals in HIV−1, SARS-CoV, and SARS-CoV2 (17,18,26-35).

In contrast to the wide-spread utilization of −1 PRF in RNA viruses, only a few cases of −1 PRF have been reported in cellular mRNAs. So far, retrovirus-like −1 PRF mechanism has been reported in the expression of three mammalian genes: including the human paternally expressed gene 10 (PEG10) (36), and the paraneoplastic antigen Ma3 & Ma5 genes (37). All of these genes are derived from retroelements and they encode viral-like proteins. In these cases, pseudoknot-dependent −1 PRF is used to express the overlapping −1 reading frame sequences in the mRNAs, with 15-30% efficiencies. Another reported case of pseudoknot-dependent −1 PRF in human mRNAs is in the C-C chemokine receptor type 5 (CCR5) mRNA (38). −1 PRF in CCR5 mRNA leads the translating ribosome to a premature termination codon in the −1 reading frame, which triggers the nonsense-mediated mRNA decay pathway to degrade the mRNA. In a computational study searching for potential −1 PRF signals in eukaryotes, it was found that up to 10% of mRNAs might utilize the −1 PRF mechanism (39). In more than 99% of the putative −1 PRF cases, a termination codon in the −1 reading frame is present shortly downstream from the frameshift site. Therefore −1 PRF in these mRNA would lead to termination of translation. It was proposed that these eukaryotic −1 PRF signals function as mRNA destabilizing elements through the nonsense-mediated mRNA decay (NMD) mechanism (38,40).

We have used bioinformatics approaches to perform a large scale search for possible cases of pseudoknot dependent −1 PRF events in human mRNAs, covering 43,191 human mRNA sequences in the NCBI RefSeq database. Several dozens of putative cases were identified. Using dual luciferase reporter assays in HEK293 cells, we confirmed that two of the putative cases harbor functional −1 PRF events. Moreover, −1 PRF in these two mRNAs differ from all of PEG10, Ma3, Ma5, and CCR5 mRNAs in that the frameshift site is located far away from the C-terminus of the normal protein and a significantly long peptide is encoded by the −1 reading frame after frameshifting.

## Materials and Methods

### Data

The Data for this study consist of 43,191 human mRNA sequences obtained from the NCBI RefSeq database (www.ncbi.nlm.nih.gov/RefSeq/).

### The pseudoknot searching program

Many algorithms and approaches had been developed to predict the formation of RNA pseudoknots. It has been proved that the general problem of predicting RNA pseudoknot is a non-deterministic polynomial time NP-complete problem. In general, most practical methods of predicting RNA pseudoknots have long running times and low accuracy for longer sequences.

To deal with the large number of long mRNA sequences, the previously reported algorithms and programs are not suitable. To facilitate our study, we have developed a new heuristic program called PKscan for the identification of RNA pseudoknots. PKscan uses a dynamic sliding window that scans through the RNA sequence, therefore there is no limitation on the length of the RNA sequence for analysis. Within each sliding window, iterating cycles of positional pairwise base matching are performed to detect complementary basepairing between two stretches of nucleotides. All possible combinations of stem and loop lengths within pre-defined ranges are interrogated for potential pseudoknot formation. PKscan can efficiently identify all potential RNA pseudoknots in any RNA sequence with unlimited length.

#### 1. Mathematical definition of relevant terms

Several relevant terms are defined mathematically as follow:

**An RNA sequence** is defined as a string of S= S_1_ S_2…_ S_n,_ where S _I_ ={A, U, G, C}.

**Legitimate basepairs** in RNA structures are the Watson-Crick basepairs and the GU wobble basepair. A legitimate basepair is represented as (S _i,_ S _j_) (i and j are positive integers and i≠ j), which must be one of the following basepairs: (A,U), (U,A), (G,C), (C,G), (G,U), or (U,G).

##### An RNA stem-loop structure

is defined by the existence of two stretches of nucleotides (separated by an intervening sequence with a certain number of nucleotides) forming complementary basepairs. An RNA stem-loop structure is mathematically represented as (S_{i},_ S_{j}_), where {i} and {i} are arrays of consecutive positive integers, and j_min_ – i_max_≥ t (j_min_ is the smallest number of the {i} array, and i_max_ is the largest number of the {i} array). The value t is a positive integer. To form a stem-loop structure, the RNA sequence folds back to itself; but the loop region must have more than t nucleotides to be physically possible. In the context of our pseudoknot detecting algorithms, t is a dynamic number depending on the stem and loop lengths of a potential pseudoknot in a particular iterating cycle of pseudoknot detection.

##### Pseudoknot

here refers to the H-type RNA pseudoknot. Mathematically, pseudoknot is defined by the simultaneous existence of two interlocking stem loop structures in a given RNA sequence (Figure 1). The two stem (double-stranded helical) regions of the stem-loop structures are represented as (S_{i},_ S_{j}_) and (S_{I’},_ S_{j’}_), where i_max_ <i’_min_+1, i’_max_ < j_min_, and j_max_ < j’_min_+3. In an H-type pseudoknot, L1 and L2 must have at least 1 and 3 nucleotide(s) respectively (L3 is optional), hence the “+1” and “+3” in the above formula.

**Figure 1:**
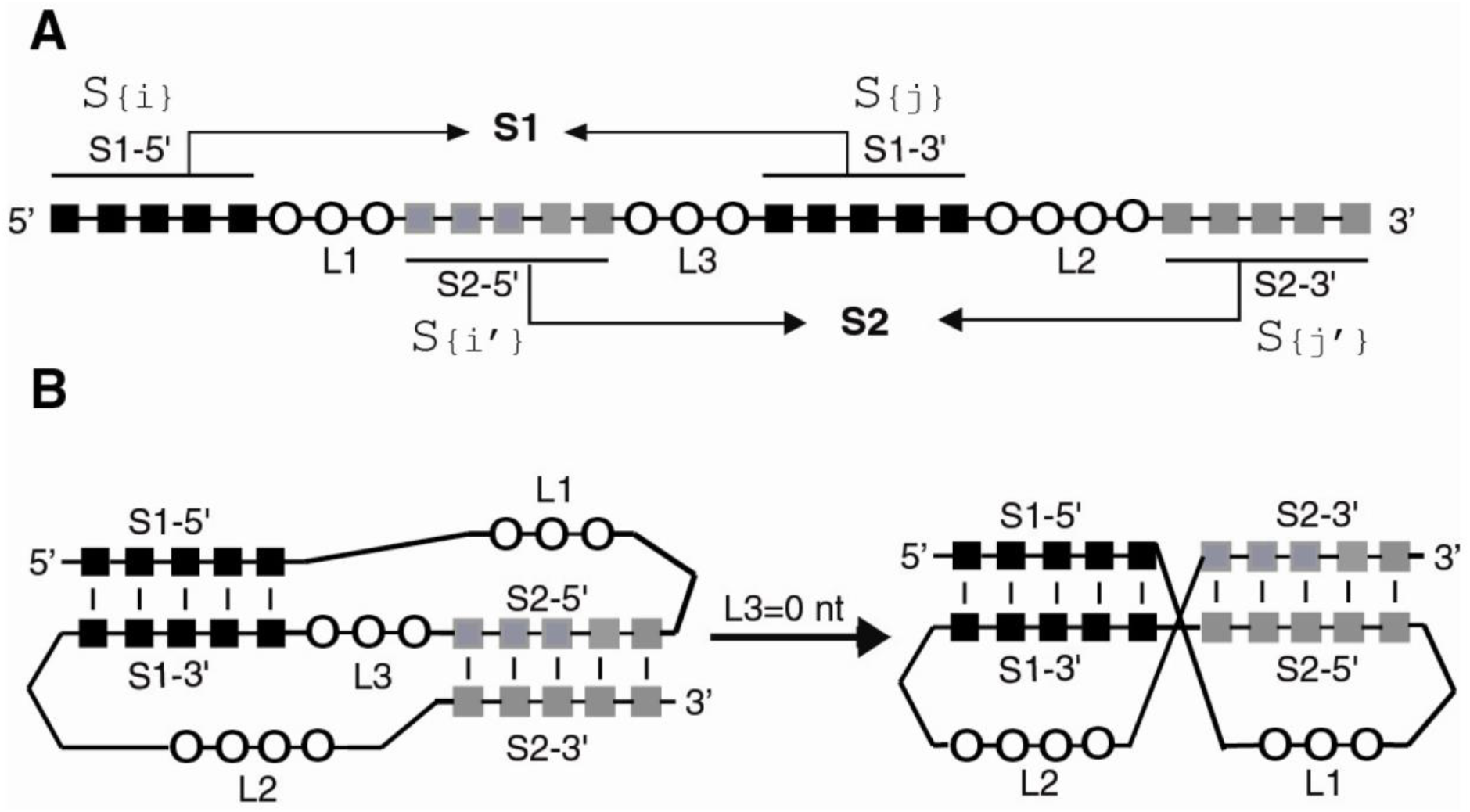
Sequence elements for forming an H-type RNA pseudoknot. Abbreviations: S1&2, stem1&2; L1-3, loop1-3. A: linear sequential arrangement of the pseudoknot-forming sequence elements. Residues involved in the formation of S1 and S2 are represented as black and gray squares respectively. Residues in the single-stranded loop region are represented as unfilled circles. B: Schematic representations of a folded pseudoknot. Left: with a non-zero L3 sequence; right: in the absence of L3, S1 and S2 can stack coaxially to form a quasi-continuous double helix. L1 and L2 locate on the same side of the double helix, with L1 crossing the major groove of S2 and L2 crossing the minor groove of S1.

(S_{i},_ S_{j}_) corresponds to the S1 stem with S_{i}_ and S_{j}_ corresponding to S1-5’ and S1-3’ stretch of nucleotide respectively. (S_{I’},_ S_{j’}_), corresponds to the S2 stem with S_{I’}_ and S_{j’}_ corresponding to S2-5’ and S2-3’ stretch of nucleotide respectively. An H-type pseudoknot must have two stems (S1 and S2) and at least two loops (L1 and L2, L3 is optional). L1 and L2 are not equivalent because L1 is crossing the major groove of the A-form RNA helical stem S2, and L2 is crossing the minor groove of the A-form RNA helical stem S1. L1 can have a minimum of a single nucleotide whereas L2 must have at least 3 nucleotides.

#### 2. Free Energy Calculation

For the involvement of pseudoknots in the −1 PRF, stability of the stimulator pseudoknot directly correlates to the ability of the pseudoknot to stimulate efficient frameshifting. It is therefore relevant and desirable to get some ideas about the relative stabilities of the identified pseudoknots in this study.

At the present time, accurate free energy calculation for RNA pseudoknots is impossible because there are no parameters available to quantify the contribution of pseudoknot specific structural features. However, the overall stability of an RNA pseudoknot is largely determined by the base paring and base stacking interactions in the stem regions, the free energy for forming the stems should provide a reasonable indicator for evaluating the relative stability of the detected pseudoknots.

Free energies of the helical stem regions of detected RNA pseudoknots are calculated based on the widely used Turner’s nearest neighbor paramerters (44).

#### 3. Algorithms of the pseudoknot searching program

The PKScan program searches for pseudoknots with pre-defined (by user) ranges for the stem and loop lengths. The default ranges are: S1 has 5 to 20 base pairs; S2 has 5 to 20 base pairs; L1 has 1 to 10 nucleotides; L2 has 3 to 50 nucleotides, and L3 has 0 to 10 nucleotides.

Within the defined ranges of stem and loop lengths, the program tests all possible combinations of stem and loop lengths to see whether the pseudoknot-forming criteria can be met. Figure 2 shows a workflow of the PKScan program, which consists of the following major steps:

**Figure 2.**
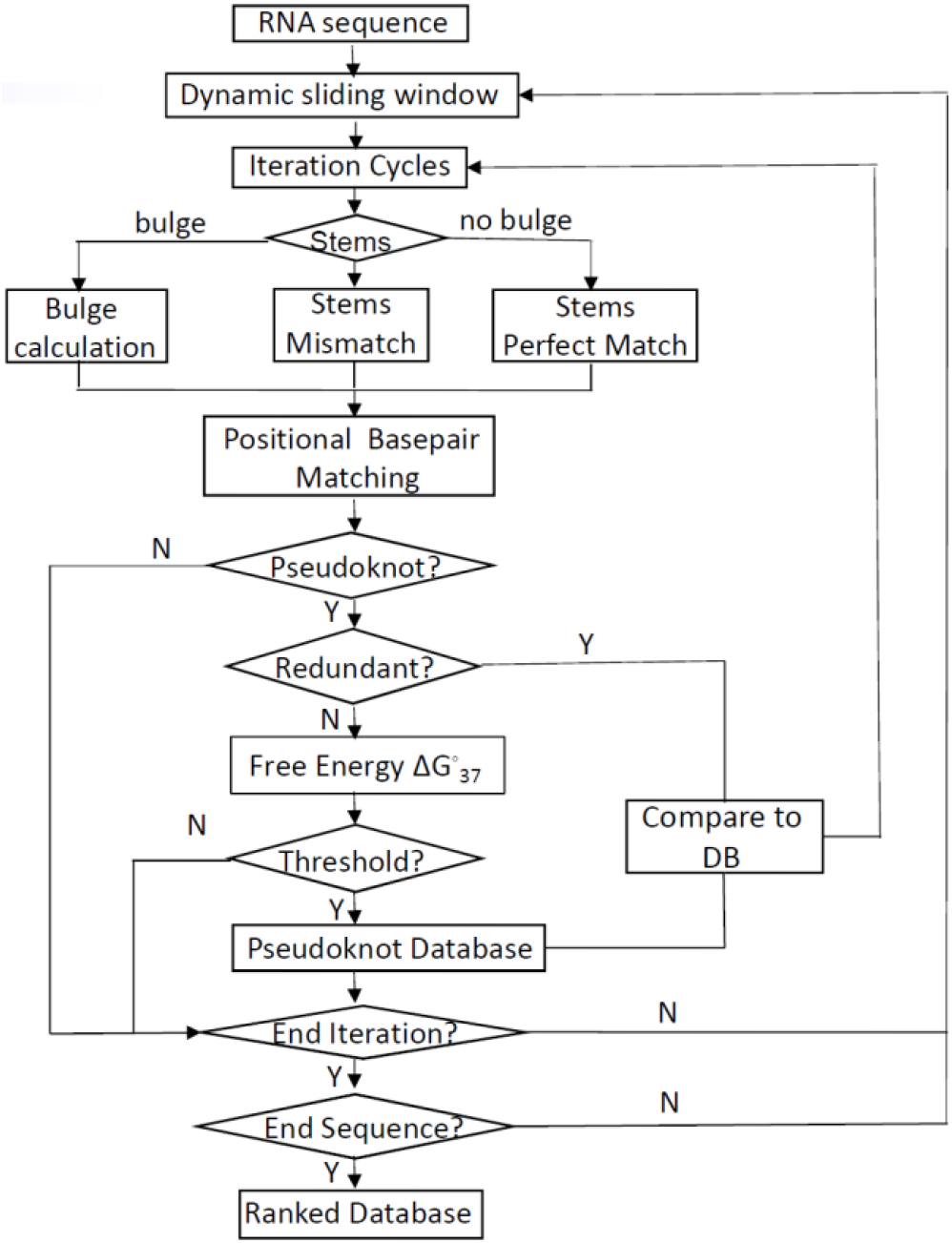
Workflow of the PKScan pseudoknot identification program

(1) Read in the RNA sequence and set the ranges of stem and loop lengths if necessary.

(2) Define the sliding window (spanning nucleotides *win[start]* to *win[end]* in the RNA sequence) as follow:

*win[start]=sn*, sn is the position of the starting nucleotide of the sliding window in the RNA sequence.

*win[end]=sn + 2 * S1[max] + L1[max] + L2[max] + L3[max] + 2 * S2[max], [max]* represents the upper limit of each of the stem and loop ranges.

Start the iterating cycles of pseudoknot detection. Each cycle has a specific combination of stem and loop lengths, i.e. specific values are calculated and assigned to s1, s2, l1, l2 and l3 (optional) as the specific lengths of the stems and loops S1, S2, L1, L2, and L3 respectively in this cycle.

(3) Depending on whether bulge or mismatchs are allowed, positional basepair matching is performed to test whether the two stems: (S_{i},_ S_{j}_) for S1, and (S_{i’},_ S_{j’}_) for S2, really exist simultaneously as complementary basepairing duplex.

For potential pseudoknots with perfectly matched stems. The sequence elements for potential pseudoknot formation are defined mathematically as follow:

~~~
Stem S1: (S_{i},_ S_{j}_)
S1-5’: S_{i}_
S1-3’: S_{j}_
{i} = [A_0,_ A_0_+s1], in increasing order.
{j} = [A_0_+ 2*s1+ s2 + l1 + l3, A0+s1+ s2 + l1 + l3], in decreasing order. Stem S2: (S_{I’},_ S_{j’}_)
S2-5’: S_{I’}_
S2-3’: S_{j’}_
{i’} = [A0+s1+ l1, A0+2*s1+ l1], in increasing order.
{j’} = [[A0+ 2*s1+ s2 + l1 + l2 + l3, A0+2*s1+ 2*s2 + l1 + l2 + l3], in decreasing order.
~~~

A_0_ is the position of the starting nucleotide of a potential pseudoknot forming sequence, s1, s2, l1, l2, and l3 are the specific lengths of stem1, stem2, loop1, loop2, loop3 of the potential pseudoknot, respectively.

Positional basepair matching is performed to evaluate whether each of the basepairs in (S_{i},_ S_{j}_) and (S_{I’},_ S_{j’}_) is a legitimate basepair of RNA. Iterating cycles of positional basepair matching are performed until all of the basepairs defined by (S_{i},_ S_{j}_) and (S_{I’},_ S_{j’}_) are tested to be true. If that turns out to be the case, then a potential pseudoknot is detected for this specific combination of stem and loop within this round of sliding window. The free energy for the stems of the pseudoknot is calculated. If any of the (S_i_, S_j_) or (S_i’_, S_j’_) basepairs is tested as false, then the iteration process is terminated, a pseudoknot cannot form with the specific combination of stem and loop lengths. The program goes back to step 3 to generate another (different) specific combination of stem and loop lengths for pseudoknot detection.

The PKscan program has the capacity to detect pseudoknots with bulges and mismatches at the stems. But in this round of study on human mRNAs, we only searched for pseudoknots with perfect matches at the stems. Therefore the algorithms for detecting pseudoknots with bulges or mismatches at the stems are not described.

(4)Use the calculated free energy value as a criterion to discard those pseudoknots with less stable stems. By default, only those pseudoknots with a free energy value lower than −18 kcal/mol are kept in the detected pseudoknot database. The default free energy threshold can be re-set by the user.

(5)Repeat steps 3-4 until all possible combinations of stem and loop lengths within the pre-defined ranges in a specific sliding window are tested for pseudoknot formation.

(6)The dynamic sliding window moves along the RNA sequence by one. A new sliding window is generated. The program goes back to step 2 for a new iterating cycle. The process keeps going until the sliding window reaches the end of the RNA sequence.

(7)All of the detected pseudoknots are ranked based on the calculated free energy values.

### mRNA sequence analysis

The mRNA sequences were subjected to analysis by a program written in C++. The search for putative pseudoknot dependent −1 PRF events in each of the mRNA sequences was carried out in several steps, as shown in the workflow in Figure 3.

**Figure 3.**
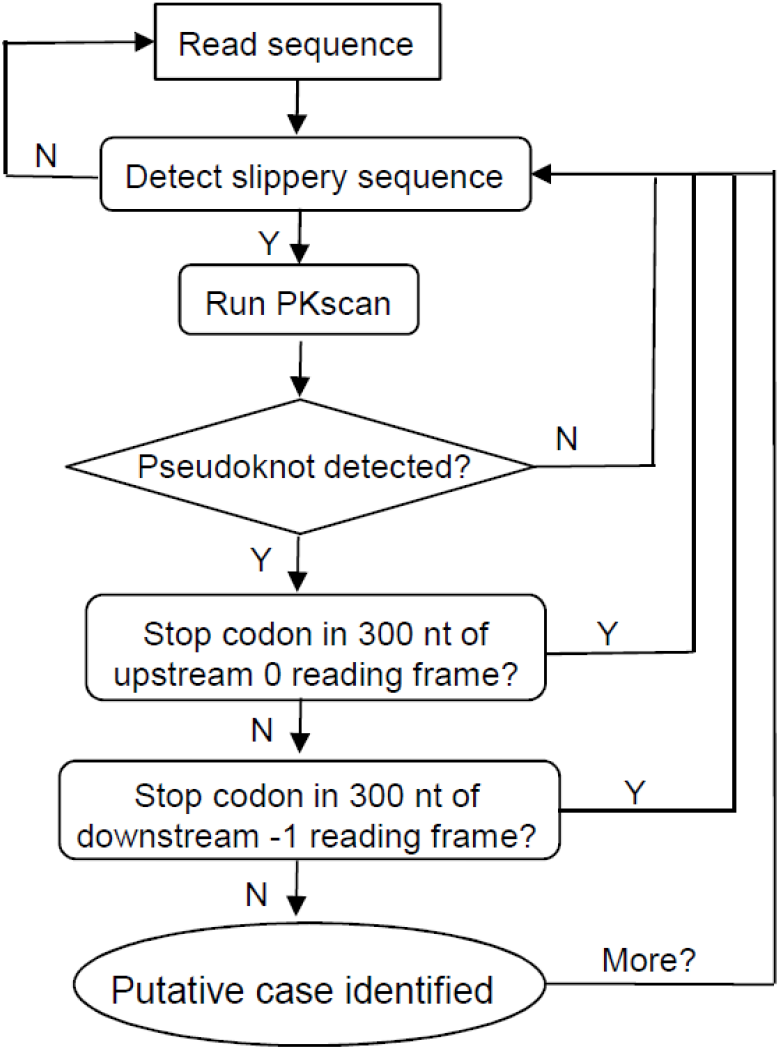
Workflow of mRNA sequence analysis.

1. Read in an mRNA sequence.
2. Scan the sequence to search for one of the following twenty heptanucleotide sequences (regardless of reading frames): AAAAAAA, AAAAAAC, AAAAAAG, AAAAAAT, AAATTTA, AAATTTT, CCCTTTA, CCCTTTT, CCCAAAA, GGGAAAC, GGGAAAT, GGGAAAG, GGGAAAA, GGGTTTA, GGGTTTT, TTTTTTA, TTTAAAC, TTTTTTG, TTTAAAT, TTTAAAG (T is used in place of U because the mRNA sequences are represented as the sense strands of the cDNAs). These heptanucleotide slippery sequences conform to the X XXY YYZ consensus and are known or suspected to be utilized *in vivo*, according to Recode V2.0: Database of translational recoding events (7).
3. For each of such putative slippery sequences identified, use PKscan to detect downstream stable pseudoknot. Due to the large number of mRNA sequences to be analyzed, we decided to restrict the pseudoknot search to only those pseudoknots with perfectly matched stems (no mismatched pair or bulge is allowed within the stems) in this round of investigation.
4. If a stable pseudoknot is detected, examine whether the 300 nucleotides upstream from the potential slippery sequence in the 0 reading frame contain any stop codon (TAG, TAA, or TGA).
5. If not, examine whether the 300 nucleotides downstream from the potential slippery sequence in the −1 reading frame contain any stop codon.
6. If not, a putative case of −1 PRF is identified. Store the case. Repeat steps 2-6 until end of the mRNA sequence.
7. Repeat steps 1-6 until all of the mRNA sequences are analysed.

### Protein Sequence Analysis

The polypeptide sequences generated by −1 ribosomal frameshifting were subjected to similarity searches using the PSI-BLAST program available at NCBI against the non-redundant protein sequences database. Protein sequence alignments were carried out by using the ClustalW program (45).

### Construction of plasmids

Plasmids for cell-based dual-Luciferase assays were constructed using a modified version of the pSGDluc plasmid (46). The original pSGDluc plasmid (obtained from Addgene) contains SV40 promotor-Renilla luciferase gene-5’ StopGo sequence-MCS-3’ StopGo sequence-Firefly luciferase gene. The MCS was replaced by a 39 bp specific sequence for ligation independent cloning (LIC), and the SV40 promotor was replaced by the CMV (human cytomegalovirus) promoter. A SmaI site is placed in the middle of the LIC sequence for vector linearization and insertion of test sequence. The modified plasmid is referred to as pSGDlucLic herein. In pSGDlucLic, transcription of the luciferase genes is driven by the strong CMV promoter, which allows constitutive and high-level gene expression in mammalian cell lines. In the mRNA, Firefly luciferase is in the −1 reading frame relative to the Renilla luciferase. Expression of the Firefly luciferase gene requires a −1 PRF event induced by the test sequence. An in-frame control plasmid, referred to as pSGDlucLic-IF, was created by inserting a basepair at the junction of the LIC sequence and firefly luciferase gene in the pSGDlucLic vector. With this insertion, the Renilla and firefly luciferase genes are transcribed in the same reading frame from the pSGDlucLic-IF vector. The 5’ and 3’ 2A StopGo sequences from the foot-and-mouth disease virus (FMDV) lead to production of the discrete Renilla luciferase and Firefly luciferase.

To construct plasmids containing the wild-type −1 PRF test sequences, the test sequence was inserted into the LIC site of the pSGDlucLic vector by ligation independent cloning (Figure 4). Each of the insert sequences was generated by a PCR reaction with Phusion DNA polymerase (Thermo Fisher Scientific) using two overlapping synthetic DNA oligonucleotides as the template and two shorter DNA oligonucleotides as the primers. Each of the primers contains a 21 nt specific LIC sequence. The PCR product was purified by agarose gel electrophoresis. The purified insert DNA was processed by T4 DNA polymerase (37°C for 45 minutes) in the presence of 25 mM dATP to generate an 18 nt 5’ overhang at both ends of the DNA insert. The pSGDlucLic vector was linearized by SmaI digestion ((37°C for 3 hours). The linearized vector was processed by T4 DNA polymerase (37°C for 45 minutes) in the presence of 25 mM dTTP to generate an 18 nt 5’ overhang at both ends. The processed insert and vector were mixed at room temperature for 20 minutes. The DNA mixture was transformed into DH5α *E. coli* competent cells to amplify the recombinant plasmids. For each wild-type test sequence, a negative control was made by changing a 0 frame codon within the slippery sequence to a stop codon; and a positive control was made by inserting one nucleotide in the slippery sequence to destroy the frameshift signal and bring the two luciferases in-frame. The controls and all of the base-pairing disruption and restoration mutants were constructed using a Gibson Assembly based mutagenesis procedure (47,48). The mutagenesis primers were used in two separate PCR reactions; The PCR products were purified by agarose gel electrophoresis; the two purified fragments were combined and mixed with equal volume of 2X NEBuilder HiFi DNA Assembly Master Mix (New England BioLabs); After incubation at 50°C for 15 minutes, the mixture was transformed into DH5α *E. coli* bacterial cells for plasmid amplification and selection. The inserted test sequences of all of the recombinant plasmids were verified by Sanger sequencing.

**Figure 4.**
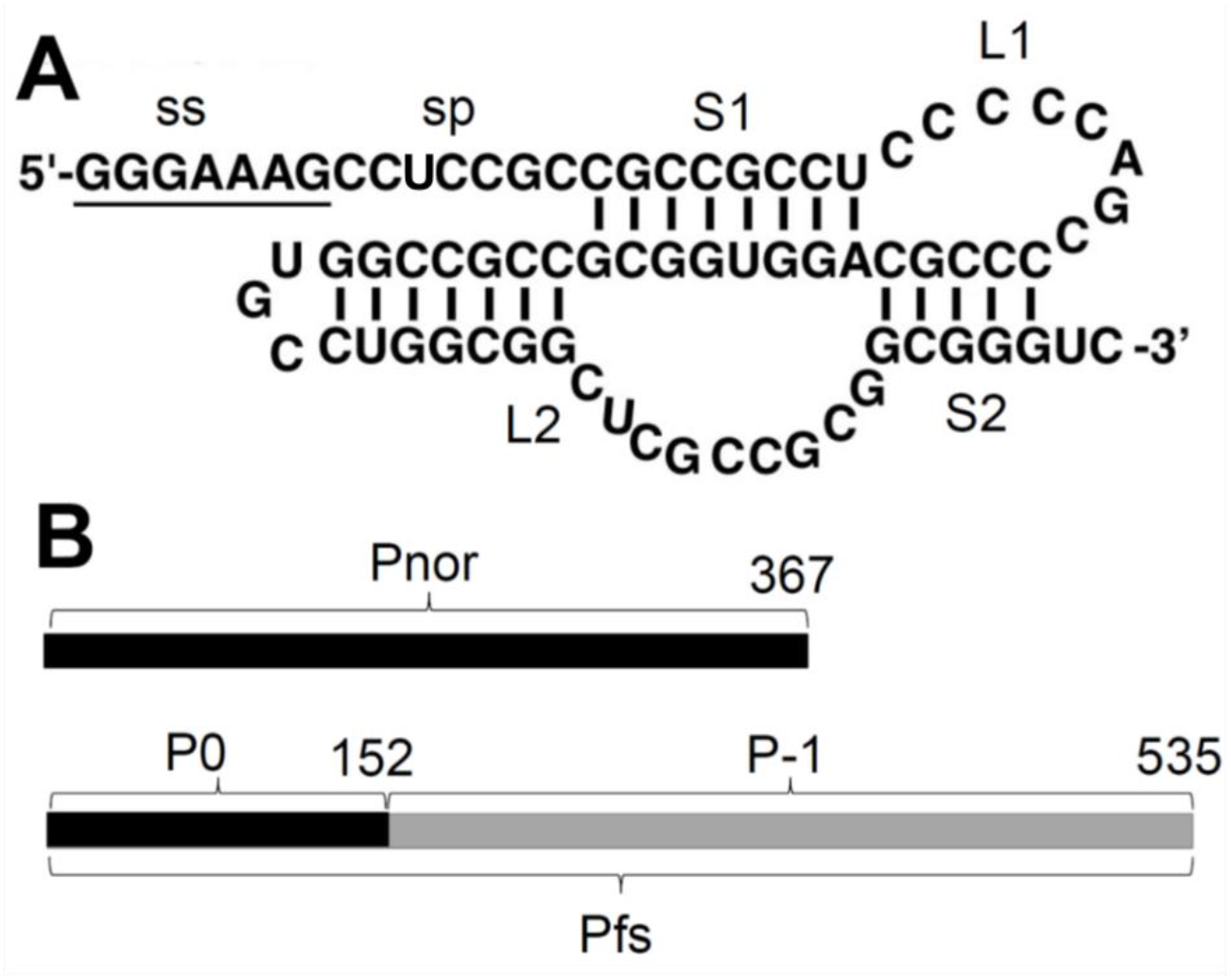
Schematic diagrams of the −1 frameshifting signals and protein products of the CDK5R2 mRNA. **A:** The cis-acting RNA signals for frameshifting: a slippery sequence (underlined) and a classical H-type RNA pseudoknot. Abbreviations used are: ss, slippery sequence; sp, spacer; S1, stem1; S2, stem2; L1, loop1; L2, loop2. Note that L2 has an extra stem-loop. **B:** The normal protein (Pnor) and the frameshift-generated protein (Pfs). Pfs is the sum of the 0 frame polypeptide produced before frameshifting (P0) and the −1 frame polypeptide produced after frameshifting (P−1). Numbers indicate the number of residues in each protein and the position of the frameshift site.

### Dual-luciferase frameshifting assays

Human embryonic kidney HEK-293FT cells (Invitrogen) were cultivated in Dulbecco’s Modified Eagle Medium (DMEM) supplemented with 10% fetal bovine serum. 24 hours before transfection, cells were harvested and resuspended in enough fresh DMEM to a final cell density of 3.5-4 ×10^5^ /mL. Seed 200µL cell suspension to each well of a 48-well culture plate. 18-24 hours after seeding, 100 ng of DNA plasmids (quantified by Qubit 3.0) were transfected into the cells in each well by Lipofectamine 2000 (Invitrogen), according to the manufacturer’s instructions. 24 hours after transfection, media were removed from the wells. 200 µl of lysis buffer was added to each well. The plate was incubated for 15 minutes at room temperature with shaking to lyse the cells. Luciferase activities were measured by using 50 µl of the lysate/sample in a 96-well plate. The GeneCopoeia Luc-Pair Firefly and Renilla Dual-Luciferase Reporter Assay System was used on a Beckman Coulter DTX 880 microplate reader. Firefly and Renilla luciferase activities were assayed subsequently in the same well. At least 9 independent transfections were carried out for each test plasmid.

−1 PRF efficiencies were calculated using the previously described method (46,49). The ratios of Firefly luciferase activity/Renilla luciferase activity (Fluc/Rluc) for the test sequences, as well as the negative and positive controls were calculated. Frameshifting efficiencies (percentage of frameshifting) were calculated as the difference between the Fluc/Rluc ratios of the test sequence and negative control divided by the ratio of the in-frame positive control.

## Results

### Strategy for identifying putative −1 PRF cases

In a previous study (37), Wills and coworkers had performed a search for putative pseudoknot-dependent −1 PRF cases in a collection of 20,763 human mRNA sequences from the RefSeq data base. The search focused on the overlapping region of the 0 and −1 reading frames to find the retrovirus-like heptanucleotide shift sites, AAAAAAC,UUUAAAC,GGGAAAC, or UUUUUUA, followed by pseudoknot detection in a 50-nt region downstream from the shift site (using the zipfold server). Only those cases in which the −1 reading frame extended at least 300 nt (encoding 100 aa) beyond the shift site were considered.

In our study, we employ a different strategy to identify putative −1 PRF cases that use a pseudoknot as the stimulator. In brief, we search for putative −1 PRF signals (a heptanucleotide shift site and a reasonably stable H-type pseudoknot 3-10 nt downstream) anywhere after 300 nt from the start codon of the 0 reading frame. Similar to the previous study, the −1 frame should have at least 300 nt beyond the shift site. Our procedure differs from the previous study in several ways: 1) much more heptanucleotide shift sites (20 vs. 4. See the Materials and Methods section for details) are considered; 2) the search for putative −1 PRF signals is not restricted in the overlapping region of the two reading frames, i.e. the frameshift sites may not locate near the end of the 0 frame; 3) pseudoknot detection is carried out by an in house developed computational method and the potential pseudoknot-forming sequences are not restricted to 50 nt (the pseudoknot can contain up to ∼150 nt using the default ranges for stems and loops, see the Materials and Methods section for details).

Using the above mentioned strategy, our search aims at identifying the following type of putative −1 PRF events: 1) occurs at one of the 20 putative heptanucleotide sequences; 2) is stimulated by an H-type pseudoknot; 3) the frameshift product contains at least 100 amino acids translated from the 0 frame and 100 amino acids translated from the −1 frame. As an example, Figure 4 shows a putative case of pseudoknot dependent −1 PRF event identified in our study. Apparently, the search does not cover all possible −1 PRF events, such as those events that are stimulated by non-pseudoknotted structures or those events that lead to termination of translation shortly after frameshifting (38,40). We also calculate the free energy of the stem regions of the pseudoknots, as a rough estimation of the stability of the pseudoknots. Pseudoknots with a calculated stem free energy higher than −18.0 kcal/mol were rejected. This cut-off value is arbitrary. Conceivably, using different cut-off value would affect the number of putative −1 PRF cases identified in the study.

### Putative −1 PRF cases

The collection of human mRNA sequences downloaded from the NCBI RefSeq database contains 43,191 sequences. A particular gene may have more than one transcript variants. After searching all of these sequences, only 30 putative −1 PRF cases are identified (transcript variants having identical frameshifting signals are counted as one case). These represent less than 0.07% of the mRNA sequences. The 30 putative cases are listed in Table 1, with relevant information about the frameshift signals and protein products. The list is organized in the order of the calculated free energy (for the stem regions) of the putative frameshift stimulator pseudoknot. It should be noted that the numbering is used merely as a convenient way of referring to specific cases; it does not represent a ranking of the putative cases based on the degree of credibility.

**Table 1.**
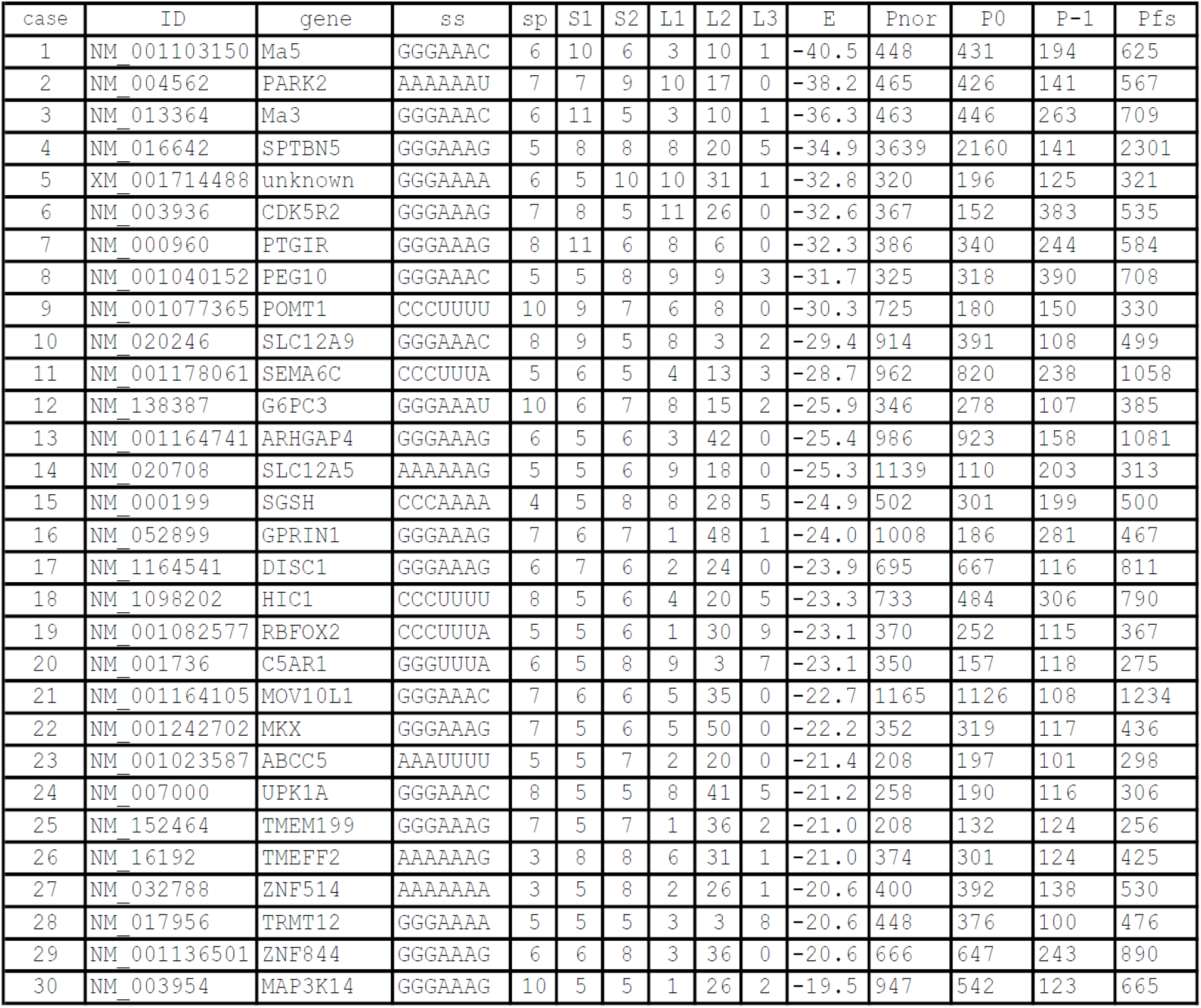
Putative cases of −1 PRF in human mRNAs. For each case, the accession ID and gene name of the mRNA are listed, together with information about the slippery sequence (ss), lengths of the spacer region (sp), the stems (S1 & S2) and loops (L1, L2, & L3) of the downstream pseudoknot. The calculated free energy of the stems of the pseudoknot is also given. The cases are listed in the order of increasing free energy values (lowest to highest). For the protein products, the lengths of the normal protein (Pnor), the 0 frame polypeptide (P0) produced before frameshifting, the −1 frame polypeptide (P-1) produced after frameshifting, and the full-length protein produced by frameshifting (Pfs) are shown. See the text and Figure 1 for more detailed description of the terms. The numbers shown as superscripts in the gene names indicate the number of transcript variants containing the same PRF signals.

As seen in Table 1, the three established or expected −1 PRF cases in human mRNAs (Ma5, Ma3, and PEG10, listed as #1, #3 and #8 respectively) are identified in our search. PEG10 mRNA has 6 transcript variants (NM_001040152, NM_001172437, NM_001172438, NM_001184961, NM_001184962, and NM_015068). Ma5 mRNA has five transcript variants (NM_001103150, NM_001103151, NM_001184924, NM_052926, NM_014550). Putative −1 PRF is detected in all of these variants (only one variant for each protein is shown in Table 1). The replication of these previously reported cases in a blind search validates the effectiveness of our approach in finding putative pseudoknot dependent −1 PRF events in a large scale computational analysis of mRNA sequences.

Figure 5 shows the putative −1 PRF signals (slippery sequence, spacer, and downstream pseudoknot) in some of the identified human mRNAs.

**Figure 5:**
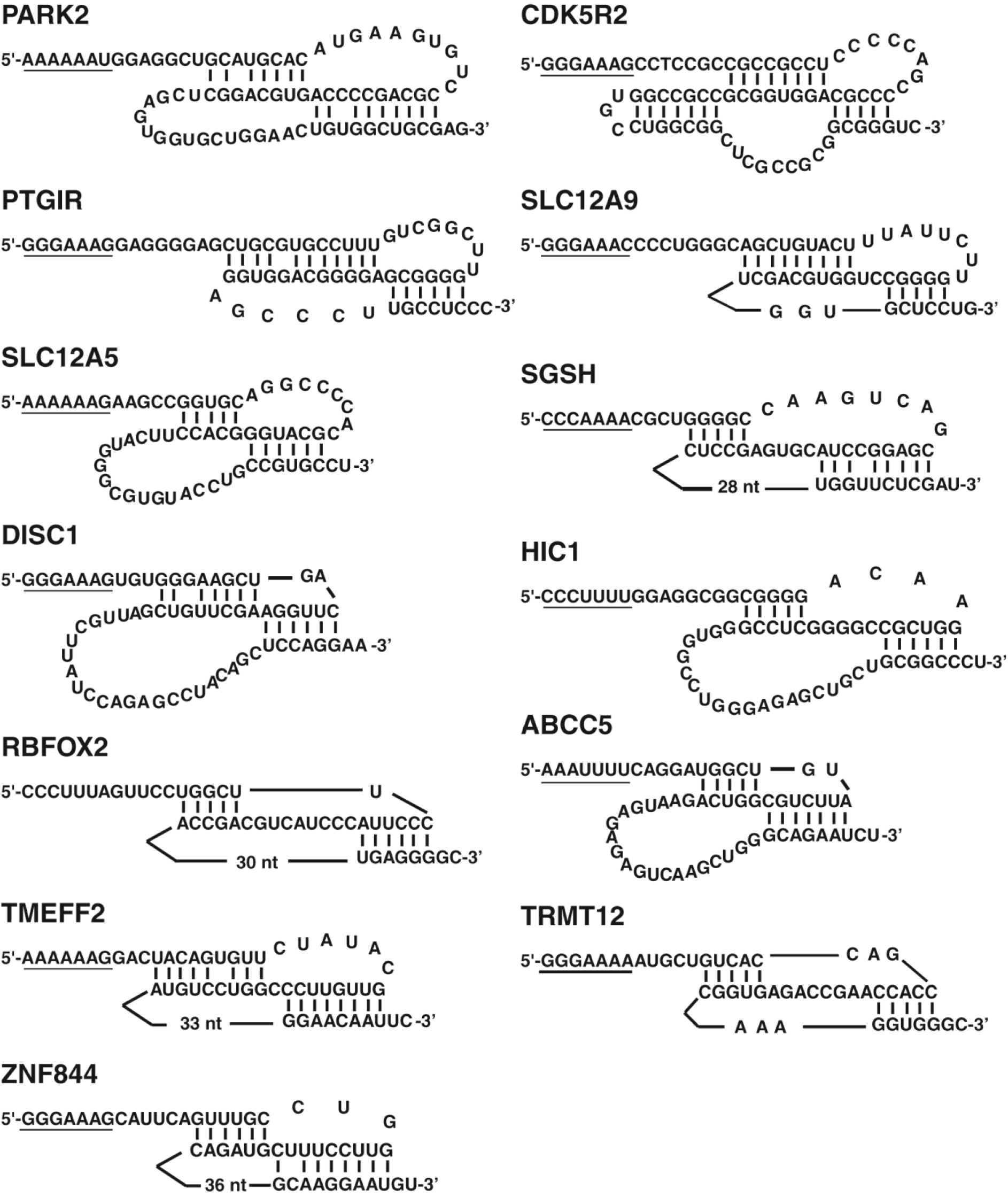
The putative cis-acting −1 PRF signals in human mRNAs. For each case, the slippery sequence (underlined), spacer, and baseparing scheme of the putative pseudoknot (or elaborated pseudoknot) are shown. See text for the full names of the abbreviations.

To get insights into the credibility of the putative −1 PRF cases identified, we performed sequence analysis on the polypeptide sequences translated from the −1 reading frame due to frameshifting. Results from the analysis provide various degrees of credibility to some of the cases, for one reason or another. Below we discuss two specific cases that are supported by bioinformatics analysis of the frameshift products and experimental evidence from dula-luciferase assays in HEK293T cells. In the following discussion, several terms are used to describe the protein products. As illustrated in Figure 4 and used in Table 1, the term normal protein (Pnor) refers to the full-length protein translated from the 0 frame without −1 frameshifting; the 0 frame polypeptide and the −1 frame polypeptide (P0 and P−1) refer to the polypeptides translated from the 0 frame before frameshifting and the −1 frame after frameshifting respectively; the protein produced by frameshifting (Pfs) refers to the full-length protein translated from the 0 and −1 frames using a −1 frameshifting mechanism. The length of Pfs is the sum of the lengths of P0 and P−1.

### −1 PRF in the CDK5R2 mRNA

Cyclin-dependent kinase 5, regulatory subunit 2 (CDK5R2) is a neuron-specific activator of CDK5 kinase. Binding of CDK5R2 to CDK5 kinase activates the enzyme (52). The slippery sequence G GGA AAG was shown to be capable of inducing −1 frameshifting in bacteria (53). The putative frameshift stimulator pseudoknot does not have a L3. Therefore the two stems S1 and S2 can potentially stack to form a quasicontinuous helix. Moreover, sequence of L2 may harbor a stem-loop structure that has the potential to stack its stem on S1 (Figure 5A). If such an elaborated pseudoknot structure really forms, it should have exceptional stability with a stack of 20 basepairs (the calculated free energy shown in Table 1 doesn’t include contribution from the extra stem S3). It is noted that the elaborated pseudoknot shown in Figure 5 for CDK5R2 mRNA is similar to the established three-stemmed frameshift stimulator pseudoknot in SARS coronavirus (54).

Human CDK5R2 normal protein (Pnor) contains 367 residues. If the putative −1 PRF event occurs, the P0 and P−1 polypeptides would contain 152 and 383 amino acids respectively, producing a frameshift protein product (Pfs) with 565 amino acids (167 amino acids longer than the normal protein). The P−1 polypeptide of CDK5R2 is the second longest among all of the −1 PRF cases identified in this study (the established case of PEG10 has the longest P−1 of 390 aa).

Using BLAST, we performed similarity searches for P−1 against the non-redundant protein sequences database. When the 383 aa sequence was used as the query, several hits were found that showed some similarities to sequence fragments of Piccolo proteins from different species. For example, a 307 aa fragment (residues 33-319) of P−1 was aligned to residues 292-588 of the protein Piccolo from Otolemur garnettii(XP_003782677) with 24% identities, 36% positives and 10% gaps. Using smaller fragments of the P−1 sequence (such as the N- & C-terminal halves) as the queries, more hits were found from a more diverse group of proteins, generally with higher degree of sequence similarity. For example, a 61 aa fragment (residues 57-117) of P−1 was matched to residues 546-606 of a protein from Drosophila virilis (XP_002056156) with 31% identities, 40% positives and 0% gaps. These sequence analysis results suggest that the P−1 polypeptide sequence of CDK5R2 is not random. It likely represents a genuine protein sequence.

To test whether the −1 PRF signals in the CDK5R2 mRNA are functional, we used dual-luciferase assays in HEK293T cells to determine the frameshifting efficiencies of a wild-type test sequence containing the −1 PRF signals and several of its mutants (Figures 6A and 6B). The test sequences are inserted between the genes for Renilla and Firefly luciferase. The Firefly luciferase gene is in the −1 reading frame relative to the Renilla luciferase gene. −1 frameshifting induced by the test sequence results in the translation of the Firefly luciferase. A negative control was constructed by inserting a stop codon in front of the test sequence. An in-frame control was constructed by inserting one nucleotide in the slippery sequence to bring the Firefly luciferase gene in frame with the Renilla luciferase gene; the slippery sequence was also scrambled.

**Figure 6.**
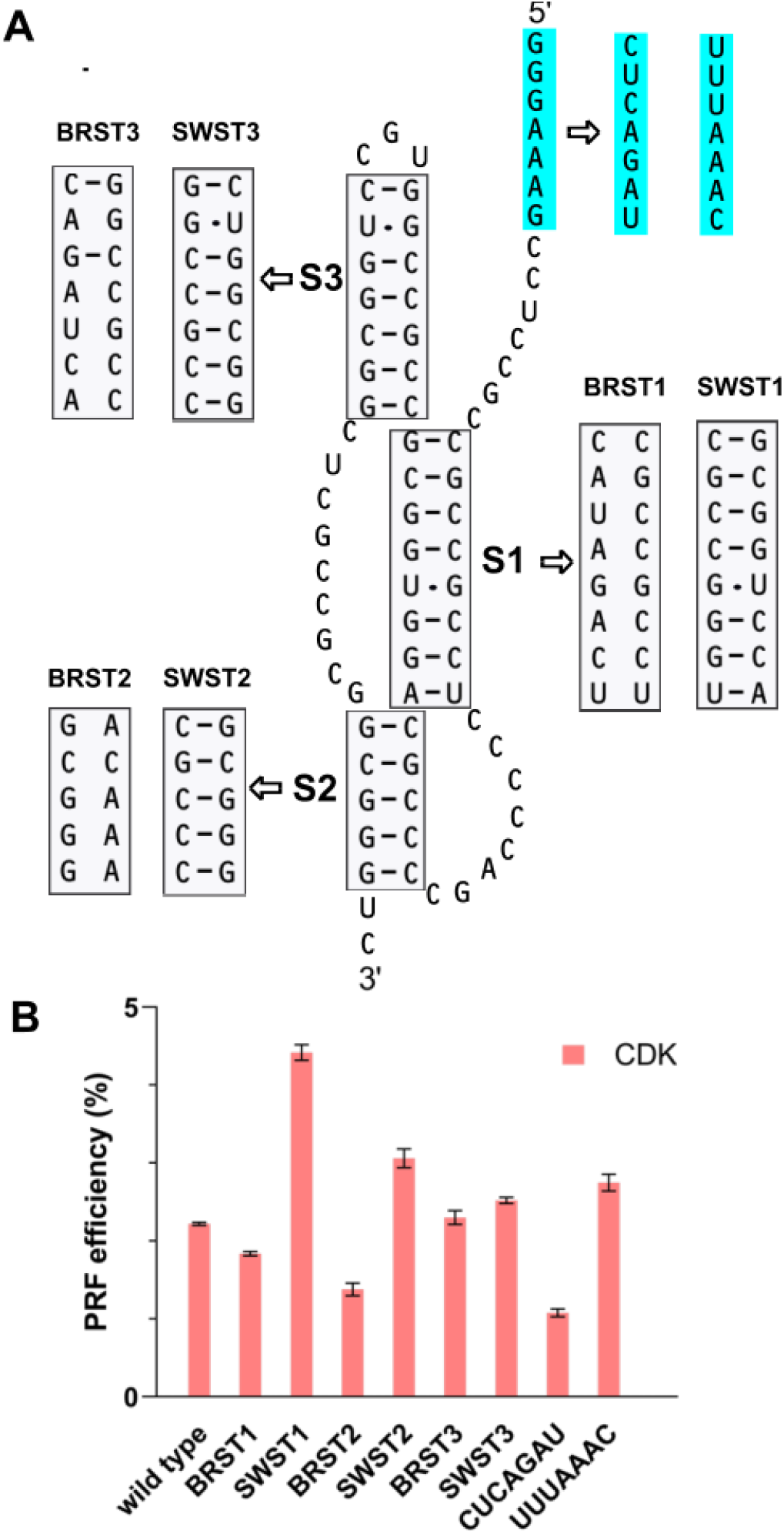
A) The −1 PRF signals in the CDK5R2 mRNA. The G GGA AAG slippery sequence is highlighted with cyan background. The downstream pseudoknot contains three stems (S1, S2, and S3). Mutations to the slippery sequence and the three stems are indicated by the open arrows. B) Functional characterization of the CDK5R2 mRNA −1 PRF signals. The −1 PRF efficiencies of the wild-type and mutant test sequences were determined by Firefly-Renilla dual-luciferase assays in HEK293T cells.

In HEK293T cells, the wild-type −1 PRF signals in the CDK5R2 mRNA stimulate efficient −1 PRF with a frameshift efficiency of 2.22 ± 0.02 %. Mutations that disrupt the slippery sequence (changed from the wild-type 5’-GGGAAAG-3’ to 5’-CUCAGAU-3’) reduced the frameshifting efficiency to 1.08 ± 0.05 %. Replacing the wild-type slippery sequence by the SARS-CoV and SARS-CoV2 slippery sequence U UUA AAC increase the frameshift efficiency to 2.75 ± 0.11 %. We had also assayed a test construct that contains the native SARS-CoV2 −1 PRF signals (the U UUA AAC slippery sequence and a downstream three-stemmed pseudoknot) using the same dual-luciferase reporter system; the frameshift efficiency was determined as 27.1 ± 0.2 %. Disruption and restoration mutations of the three stems all affect the frameshift efficiency (Figure 6B). While the stem1 disruption construct reduces the frameshift efficiency, the stem1 restoration construct doubles the frameshift efficiency. In this stem1 restoration construct, the two strands of nucleotides forming the stem are swapped from the wild-type and the 5’ strand now becomes purine-rich. It is noticed that in most of the naturally occurring pseudoknots that stimulate −1 PRF with a high efficiency, the 5’ stem1-forming strand is purine rich. The stem2 disruption construct reduces the frameshift efficiency significantly (∼40% reduction), while the stem2 restoration construct increases the frameshift efficiency by about 40%. The stem disruption and restoration constructs of stem3 both increase the frameshift efficiency. These data show that the −1 PRF signals in the CDK5R2 mRNA are functional in human cells, and the predicted pseudoknot functions as a frameshift stimulator.

### −1 PRF in the SEMA6C mRNA

Human SEMA6C has three variants in the NCBI Reference Sequence database (962, 930, and 922 aa for variant 1, 2 and 3 respectively). Compared to variant 1, the mRNAs of variants 2 and 3 apparently lack an exon encoding 32 aa and 40 aa respectively. The three variants otherwise have identical protein and DNA sequences.

Figure 7A shows the putative −1 PRF signals in the SEMA6C mRNA. The slippery sequence of C CCU UUA is identical to the established slippery sequence utilized by giardiavirus (GLV) at the viral *gag-pol* junction for −1 frameshifting (43). A spacer sequence of five nucleotides separates the slippery sequence and a downstream pseudoknot. A −1 ribosomal frameshifting event induced by these signals would produce a polypeptide of 238 amino acids encoding by the −1 reading frame. The normal protein sequence contains 142 amino acids residues after the frameshifting site (Table 1).

**Figure 7.**
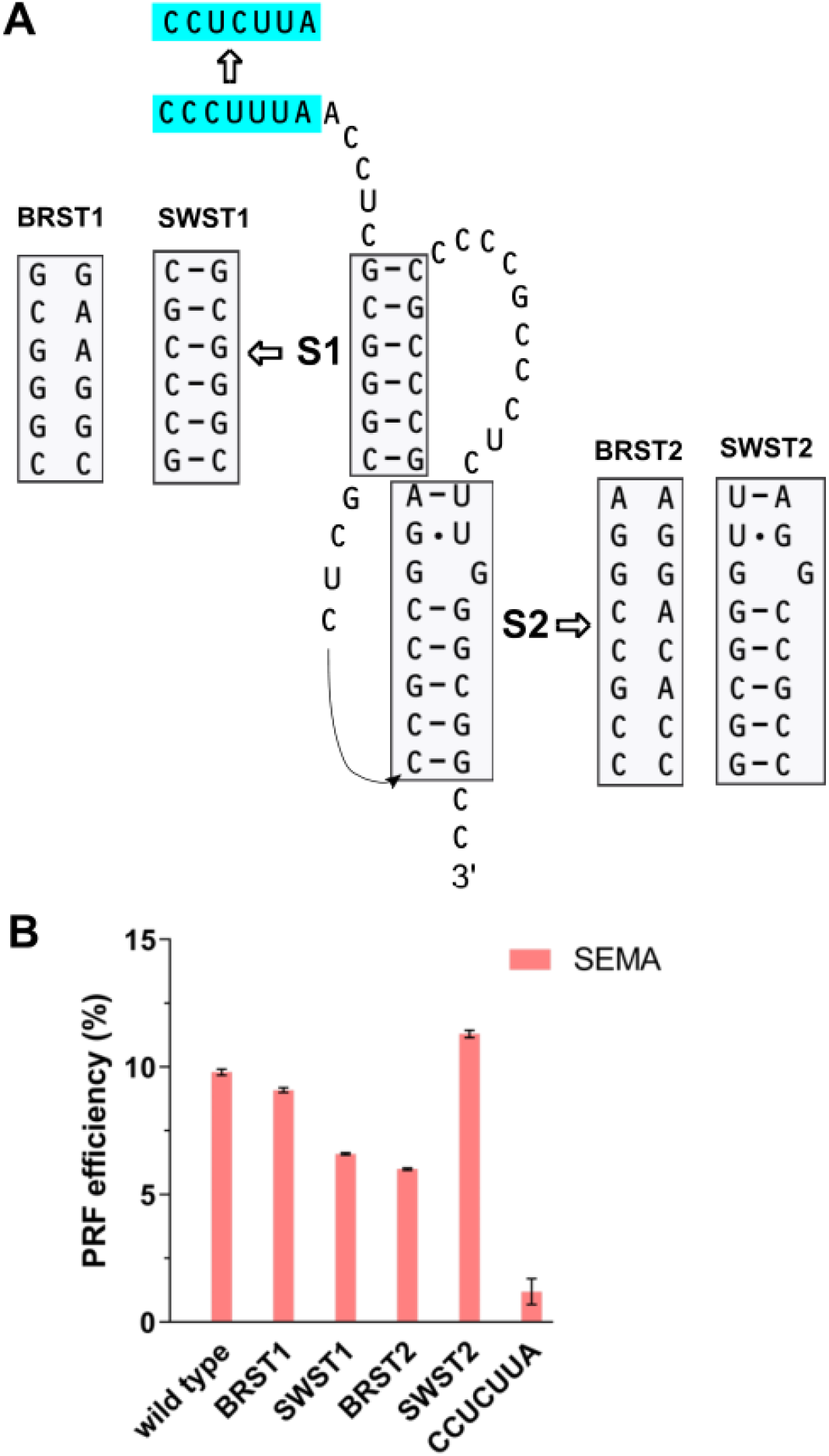
A) The −1 PRF signals in the SEMA6C mRNA. The C CCU UUA slippery sequence is highlighted with cyan background. The downstream pseudoknot contains two stems (S1 and S2). Mutations to the slippery sequence and the two stems are indicated by the open arrows. B) Functional characterization of the SEMA6C mRNA −1 PRF signals. The −1 PRF efficiencies of the wild-type and mutant test sequences were determined by Firefly-Renilla dual-luciferase assays in HEK293T cells.

We performed a similarity search for the polypeptide generated by −1 PRF using BLAST against the non-redundant protein sequences database. When using the 238 amino acids sequence as the query, only one hit was found: AAI14624, which is annotated as a human SEMA6C protein with 537 residues. Residues 74-238 of the query are identical to the last 165 residues (aa 373-537) of AAI14624 except at one residue (Ala-137 vs Val-446). More interestingly, the protein AAI14624 is encoded by a complete cDNA clone (IMAGE:40036285), indicating that the protein is actually expressed. AAI14624 contains 537 residues; residues 1-371 are identical to corresponding sequence of SEMA6C variant 1 and residues 373-537 match the C-terminal 165 residues of the frameshift product. Therefore, AAI14624 likely represents a novel alternatively spliced isoform of human SEMA6C with the N-terminal ∼70% of the sequence being encoded by the zero reading frame and the C-terminal ∼30% of the sequence being encoded by the −1 reading frame (using the mRNA of variant 1 as a reference). We also used the residues 1-73 of the frameshift product as the query in a BLAST search. Only one hit was found: AES06293, which is annotated as a partial sequence of a SEMA6C protein from Mustela putorius furo. Residues 15-60 of the query show a high degree of similarity to the last 46 residues of AES06293 (identities 54%, positives 63%, no gap). From the above sequence analysis, it is clear that ∼90% of the 238 amino acids residues of the frameshift product match to SEMA6C protein sequences in the database. These results provide strong support for the utilization of −1 ribosomal frameshifting mechanism in the translation of human SEMA6C mRNAs. A SEMA6C protein isoform produced by −1 ribosomal frameshifting would contain 1058 amino acids residues.

Using dual-luciferase assays in HEK293T cells, we showed that the wild-type −1 PRF signals in the SEMA6C mRNA stimulate efficient −1 PRF with a frameshift efficiency of 9.8 ± 0.12%. Mutations that disrupt the slippery sequence (changed from the wild-type 5’-CC<u>CU</u>UUA-3’ to 5’-CC<u>UC</u>UUA-3’, by swapping the two underlined nucleotides) greatly reduced the frameshifting efficiency to 1.2 ± 0.5 % (Figure 7B). The stem1 disruption construct reduces the frameshift efficiency slightly. We examined the sequence this particular designed construct and found that an alternative stem 1 (with a 5’-GGGC-3’ and 3’-CCCG-5’ base-paring scheme) could form, which might explain the corresponding frameshift efficiency. The stem1 restoration construct reduces the frameshift efficiency by about 33% to 6.6 ± 0.05 %. In this stem1 restoration construct, the two strands of nucleotides forming the stem are swapped from the wild-type. The 5’ stem1-forming strand in the wild-type is purine rich, and it becomes pyrimidine rich in the stem1 restoration construct. This may explain the reduction in frameshift efficiency of the stem1 restoration construct, based on the same reasoning as in the CDK5R2 case. The stem2 disruption construct reduces the frameshift efficiency by almost 40%, while the stem2 restoration construct increases the frameshift efficiency by about 15% to 11.3 ± 0.14 %. These data show that the −1 PRF signals in the SEMA6C mRNA are functional in human cells, and the frameshifting event is stimulated by a downstream pseudoknot.

## Discussion

We have performed a large-scale search for putative pseudoknot-dependent −1 PRF events in human mRNAs. Only 30 putative cases were identified, accounting for less than 0.07% of the mRNA sequences. Among these 30 identified cases, three cases are previously reported for PEG10, Ma3 and Ma5 mRNAs. Replication of these cases proves that our approach is effective.

The putative −1 PRF cases differ from each other in terms of the cis-acting mRNA signals for frameshifting and the parameters of the protein products. For one reason or another, some of the cases may seem more credible than the others. Very likely, some of the cases are just results by coincidence and don’t represent real −1 PRF events. We note that in a number of cases, the sequence of the P−1 polypeptide is compositionally biased and has no similarity to known protein sequences in the database (as detected by BLAST search). Presumably, these cases have a bigger chance of being false positives.

Unlike the three established −1 PRF cases in PEG10, Ma3, and Ma5, the genes in the other identified putative cases do not have retroviral origins and they do not encode virus-like proteins. Moreover, in all but four (ABCC5, DISC1, PTGIR and ZNF844) of the cases, the frameshift sites locate hundreds of nucleotides upstream from the stop codon of the 0 reading frame, i.e. a C-terminal sequence of significant length in the normal protein is replaced by the P−1 polypeptide sequence in the frameshift protein product Pfs. In the cases of CDK5R2, SLC12A9 and SLC12A5, the frameshift sites even locate within the 5’-half of the protein coding sequence (Table 1), which means that less than half of the normal protein sequence is present in the frameshift protein product Pfs. In a typical viral or viral-like −1 PRF event, the frameshift site locates within tens of nucleotides from the 0 frame stop codon; the frameshift protein product contains almost the entire sequence of the normal protein, plus a C-terminal polypeptide sequence encoded by the −1 frame. Apparently, some of the putative −1 PRF cases identified in our study are very different from those in viral or viral-like mRNAs. To the best of our knowledge, there is no established case of functional −1 PRF event with a frameshift site locating far upstream from the 0 frame stop codon. Our study identifies many such putative PRF cases. Validation of the −1 PRF signals in the CDK5R2 and SEMA6C mRNAs by experimental procedures indicates that some of the cases identified by the computational study are functional. These results expand the repertoire of the −1 PRF phenomenon and the protein*-*coding capacity of the human transcriptome.

As detailed in the Materials and Methods section, we have restricted our search for putative −1 PRF events in this study to those cases that: 1) are stimulated by a downstream H-type pseudoknot with certain stem and loop lengths, as well as decent thermal stability (measured by the calculated free energy of the stems); 2) contain at least 100 aa residues in both the P0 and P−1 polypeptides. The search therefore does not intend to be exhaustive. It should also be noted that the pseudoknot searching program used in this study could only detect pseudoknots with either perfect stems or stems containing only a single base pair mismatch (see Figure 5 for some examples). Considering these restrictions and limitations, we expect that human mRNAs should contain more functional −1 PRF events than currently known.

## Acknowledgements

The work was supported by the start-up fund and a seed grant from Southern Illinois University Carbondale to Z.D.

